# The Cdc42 GTPase activating protein Rga6 promotes the cortical localization of Septin

**DOI:** 10.1101/2021.03.30.437639

**Authors:** Shengnan Zheng, Biyu Zheng, Zhenbang Liu, Wenfan Wei, Chuanhai Fu

## Abstract

Septins are a family of filament-forming GTP-binding proteins that regulate fundamental cellular activities such as cytokinesis, cell polarity, and membrane remodelling. In general, Septin filaments function as barriers and scaffolds on the cell cortex. However, little is known about the mechanism that governs the recruitment and localization of the Septin complex to the cell cortex. Here, we identified the Cdc42 GTPase activating protein Rga6 as a key protein involved in promoting the localization of the Septin complex to the cell cortex in the fission yeast *Schizosaccharomyces pombe*. Rga6 interacts with the Septin complex and colocalizes with the Septin complex on the cell cortex. Live-cell microscopic analysis further showed Septin enrichment at the cortical regions adjacent to the growing cell tip. The Septin enrichment likely plays a crucial role in confining active Cdc42 to the growing cell tip. Hence, our findings support a model that Rga6 regulates polarized cell growth partly through promoting targeted localization of the Septin complex on the cell cortex.

## INTRODUCTION

Septins are evolutionarily conserved GTPases and mainly localizes to membrane structures within the cell (1–4). On the cortical membrane, Septins form high-order structures, including short filaments, rings, and gauzes, in a cell-cycle and/or location-dependent manner (1, 4–7). These Septin high-order structures could function as scaffolds responsible for recruiting a wide range of proteins to regulate cytokinesis and/or as diffusion barriers to compartmentalize membranes (3, 4, 8–11). In addition, Septins play critical roles in membrane remodelling and in regulating cortical rigidity (3, 12–16). Despite significant progress in understanding Septin functions, how Septins are localized to the cortical membrane remains elusive.

The membrane affinity of Septins may be dictated by their intrinsic property since an N-terminal polybasic region is present in most Septins (3, 17, 18). Consistently, recombinant Septins can bind silica beads decorated with phospholipids, dependent of Septin concentration and bead size (19). Moreover, *in vitro* lipid-binding assays established that Septins bind phosphoinositides (6, 17, 18, 20). The fission yeast *Schizosaccharomyces pombe* has seven Septins, i.e. Spn1-7 (20). Spn1-4 function in vegetative cells while Spn2 and Spn5-7 mediate forespore membrane extension during sporulation (20, 21). Similarly, the N-terminal region of Spn1-4 and Spn7 contains a cluster of basic amino acids, and *in vitro* phosphoinositide binding assays with Spn2 and Spn7 showed that both Spn2 and Spn7 have an affinity for PtdIns(4)P and PtdIns(5)P (20). Recent evidence suggests that Septins are sensors of micron-scale membrane curvature and are often enriched at the membrane regions with high curvature, such as the yeast bud neck, the bases of membrane branches, and the cell projection tips (19, 22). Nevertheless, Septins can also localize to the cortical regions with low curvature. For instance, during interphase, fission yeast Septins localize as short filaments on the cell sides and form complex high-order structures through annealing and end polymerization (5). How Septins are localized to the cell cortex with low curvature remains poorly understood, and the function of the cortically localized Septins is elusive.

In the budding yeast *Saccharomyces cerevisiae*, Septins are recruited to the cortical region of the budding site by the Rho GTPase Cdc42, the master regulator of cell polarity, via the effector proteins of Cdc42, i.e. Gic1 and Gic2 (23–25). The cortically localized Septins subsequently recruit Cdc42 GTPase activating protein (GAP) to inhibit Cdc42 activity and the inhibitory effect on Cdc42 is counteracted by polarized exocytosis (26). These feedback controls among Cdc42, Septins, and exocytosis then promote the formation of the Septin ring at the bud neck (26). Therefore, the interplay between Septins and Cdc42 GAP proteins play critical roles not only in Septin ring formation but also in polarized cell growth. Interestingly, similar to Septins, most RhoGAP proteins contain membrane-binding domains such as BAR, PH, and PBR (polybasic region) (27). Whether the interplay between RhoGAP proteins and Septins is important for their localization to the cell cortex is an interesting question to pursue.

Nine RhoGAP proteins exist in the fission yeast (28). Among them, Rga3, Rga4 and Rga6 localize to the cell cortex and have GAP activity for Cdc42 (29–32). Functional characterization shows that Rga4 and Rga6 work in concert to regulate cell polarity by restricting active Cdc42 to the growing cell tip (29).

In this study, we identified Rga6 as an interacting protein of the Septin complex and demonstrated that the interaction between Septins and Rga6 promotes the cortical localization of Septins. On the cell cortex, Rga6 and Septins synergize to regulate cell growth by confining active Cdc42 to the cell tips. Hence, this present work establishes a role of Septins in regulating the targeted localization of active Cdc42 required for polarized cell growth.

## MATERIALS AND METHODS

### Yeast genetics

Yeast strains were created either by the random spore digestion method or by tetra-dissection analysis. Gene deletion and tagging were performed by the PCR-based method using the pFA6a series of plasmids, and yeast transformation was carried out by the lithium acetate method (33). The strains used in this study are listed in Supplementary Table S1. For live-cell imaging, the strains were cultured in Edinburgh minimal media supplemented with Adenine, Leucine, Uracil, Histidine, and Lysine (ForMedium, Hunstanton, Norfolk, United Kingdom). For biochemistry, cells were cultured in yeast extract (YE) media containing the five supplements indicated above.

### Molecular cloning

Plasmids used in this study are listed in Supplementary Table S2. The restriction enzymes used for molecular cloning were purchased from NEB (www.neb.com), and cloning was carried out by the conventional digestion/ligation method or by ClonExpress II One Step Cloning (www.vazymebiotech.com).

For generating the Septin constructs, the four *septin* genes were first amplified by PCR with a yeast cDNA library. For generating the pETDuet plasmid carrying Spn1 and Spn2-GST (pCF.3623), *spn1* was digested with BamHI and NotI and spn2 was digested with BglII and XhoI. Both digested products were then cleaned with a PCR purification column (TIANGEN), and ligation was then performed using the T4 DNA ligase (NEB). GST tag was amplified by PCR and was inserted into the pETDuet plasmid, downstream of *spn2*, by the recombination method with a One-step cloning kit (ClonExpress II One Step Cloning Kit).

For generating pACYCDuet carrying Spn3 and Spn4 (pCF.3301), *spn3* was digested with PstI and NotI and *spn4* was digested with BglII and XhoI. Ligation was then performed using the T4 DNA ligase (NEB).

For generating pFastBac-Rga6-GFP-His (pCF.3805), *rga6* was amplified from pCF.3080 and digested with BssHII and NotI for further ligation.

For generating rga6 full-length and deletion-truncation mutant plasmids, the inserts obtained by PCR were digested with BglII and NotI and ligated into a pJK148 vector with a T4 DNA ligase (NEB).

### Live-cell microscopy and data analysis

Live-cell imaging was performed with a PerkinElmer Ultraview spinning disk confocal microscope equipped with a Nikon Apochromat TIRF 100X 1.49NA objective and a Hamamatasu C9100-23B EMCCD camera, following the method as described previously (34). Briefly, cells in the exponential phase were collected and sandwiched between a EMM (with supplements: Adenine, Leucine, Uracil, Histidine, and Lysine) agar pad and a coverslip. Images were acquired with Volocity (PerkinElmer, Waltham, Massachusetts, USA) at room temperature. Unless otherwise specified, image stacks containing 11 planes were taken with a 0.5 μm step size and the time-lapse images were taken every 2 minutes (500 ms exposure for GFP/mNeonGreen and tdTomato).

For the FRAP experiments and the colocalization of Rga6-3GFP and Spn1-tdTomato, imaging was performed with a ZEISS LSM880 (Airyscan) confocal microscope equipped with a ZEISS flat field achromatic100X 1.4NA objective and a laser scanner. Briefly, images were acquired with ZEN black software; bleaching was conducted using 15% 543 nm laser power (10 repeats), and the recovery time-lapse images were taken every 60 seconds at the middle plane using 3% 543 nm laser power. The FRAP data were exported from the ZEN black software.

Images were analyzed and measurements were performed, using MetaMorph 7.7 (Molecular Devices, San Jose, CA, USA) and ImageJ 1.52 (NIH). Graphs were generated and statistical analysis was performed, using Kaleidagraph 4.5 (Synergy Software,Reading, PA, USA).

### Proteins purification and Size-exclusion chromatography (SEC)

For producing the Septin complex with a GST tag and GST protein, BL21(DE3) *Escherichia coli* cells were transformed with the duet Septin expression plasmids, selected with ampicillin and chloramphenicol, or the plasmid pGEX-6p-1, and induced to express with 1 mM IPTG at an OD_600_ of 0.8-1.0. After 24 hours of growth at 22°C, cells were harvested by centrifugation at 5,000 g (relative centrifugal force) for 15 min. Cell pellets were resuspended in lysis buffer (50mM Tris (pH 8.0), 300mM NaCl) and crushed for 2 minutes with a high-pressure crusher (700-800 pressure), and the cell extract was centrifuged at 4°C for 40 minutes at 13,000 rpm. The supernatant was then incubated with 500 μl Glutathione Sepharose resins (17-5279-02, GE Healthcare Bio-Science) for 90 minutes. The resins were washed with GST wash buffer (50 mM Tris (pH 8.0), 300 mM NaCl, and 0.1% Triton X-100) (10X column volume) and were kept in storage buffer (50 mM Tris (pH 8.0), 300 mM NaCl, 0.1% Triton X-100, and 20% Glycerol) at 20°C.

For producing Rga6-GFP, Sf9 insect cells were transfected with the bacmid pFast-Rga6-GFP-His. After 3 rounds of virus infection, cells were harvested by centrifugation at 5,000 g for 10 min. Cell pellets were lysed in lysis buffer (50mM NaH2PO4 (pH 8.0), 300 mM NaCl, and 30 mM imidazole) and crushed for 2 minutes with a high-pressure crusher (200-300 pressure), and the cell extract was centrifuged for 20 minutes at 13,000 rpm at 4 °C. The supernatant was then incubated with 2 ml Ni-NTA Superflow resins (Cat No.30430, QIAGEN) for 30 minutes. The resins were washed with lysis buffer (50 mM NaH2PO4 (pH 8.0),300 mM NaCl, and 30 mM imidazole) (10X column volume), and bound proteins were eluted with elution buffer (50 mM NaH2PO4 (pH 8.0), 300 mM NaCl, and 300 mM imidazole).

For producing recombinant proteins Spn2-BFP or BFP proteins, BL21(DE3) *Escherichia coli* cells were transformed with the duet expression plasmid carrying Spn2-BFP or pET28a-BFP plasmid and induced to express with 1 mM IPTG at an OD_600_ of 0.8-1.0. The similar procedure as above, used for purifying Rga6-GFP, was then used for the purification of Spn2-BFP or BFP. The purified recombinant proteins Spn2-BFP or BFP were concentrated to 2 ml and were further purified by SEC with a HiLoadTM 16/600 superdexTM 200 pg column (28-9893-35; GE Healthcare) in SEC buffer (50 mM Tris (pH 7.5), 150 mM NaCl, 2 mM MgCl_2_, and 1 mM DTT).

### Co-Immunocoprecipitation (co-IP) assay

Co-immunoprecipitation assays were performed with Dynabeads protein G beads (10004D; Thermo Fisher) and strains expressing the indicated GFP and tdTomato proteins in TBS buffer containing 0.1% Triton X-100. Cell lysates were prepared by grinding in liquid nitrogen with a mortar grinder RM 200, and dissolved in TBS lysis buffer containing 0.1% Triton X-100 and cocktail protease inhibitors. Dynabeads Protein G beads bound with the GFP antibody were incubated with the cell lysates for 1 hour at 4 °C, followed by washing with 1x TBS buffer containing 0.1% Triton X-100 for five times and with 1x TBS buffer for one time. Immunoprecipitated proteins were then analyzed by western blotting with antibodies against GFP (600-101-215; Rockland) and tdTomato (Home-made; GenScript).

### Analysis of protein expression

For analysis of protein expression levels, proteins extracts were prepared by the TCA lysis method. Exponential cells were collected from 15OD culture. After washed 1 time with 1ml distilled deionized water (ddwater), the cells were resuspended with 50 μl 20% TCA and vortexed. About 200μl glass beads was added, and cells were disrupted, using Retsch MM400 for 4-5 minutes. Vortex was performed after 50 μl 20% TCA was added, and vortex was performed again after 400μl 5% TCA was added. After removing the glass beads, samples were centrifuged at 13,000 rpm for 10min at 4°C to remove the supernatant. Finally, the precipitate was resuspended with 100μl 1X SDS sample buffer and 25μl 1.5M Tris (pH=8.0), followed by boiling at 100°C for 5 minutes. The proteins were analyzed by western blotting.

### GST pull-down assay

For the GST pull-down assay with only recombinant proteins, recombinant proteins Rga6-GFP and GFP were incubated with 20 μl glutathione resins bound with GST-fused proteins or with empty glutathione resins in 200 μl TBS buffer (50mM Tris (pH=7.5), and 150 mM NaCl) supplemented with 0.1 % Triton X-100 and 40 mM imidazole. After incubation of 90 minutes at 4°C, the resins were washed 5 times with TBS+0.1% Triton X-100 and 1 time with TBS. The resins were boiled in SDS-PAGE sample buffer for 5 minutes, and protein samples were analyzed by SDS-PAGE and immunoblot with antibodies against GFP and GST.

For testing the interaction of recombinant GST-fused proteins with Rga6-GFP, cell lysis containing Rga6-GFP was prepared and used for incubation with the indicated recombinant GST-fused proteins. After incubation of 90 minutes at 4°C, the GST resins were washed 5 times with TBS+0.1% Triton X-100 and 1 time with TBS. The resins were boiled in SDS-PAGE sample buffer for 5 minutes, and protein samples were analyzed by SDS-PAGE and immunoblot with antibodies against GFP and GST.

### Liposome reconstitution assay

The binding between recombinant proteins and lipids was evaluated using a lipid composition of 90% PC (egg, chicken; 840051; Avanti Polar Lipids) and 10% PS (Brain,840032p;Avanti Polar lipids) (35). Lipids were mixed in chloroform solvent (90 μl PC+10 μl PS, all at 10 mg/ml), dried by a steam of nitrogen for 5 minutes, and dissolved with 500 μl mineral oil (330760; Sigma) (36). The mixture was vortexed for 5 minutes, and 30 μl recombinant proteins at the indicated concentration, pre-incubated on ice for 20 mintues (in the SEC buffer), were added to the lipid-mineral mixture (100μl). The protein-containing mixture was then vortexed again for 30-60s and a drop of the lipid-protein mixture was added to a glass profusion chamber for imaging with a spinning-disk confocal microscope.

## RESULTS

### Microscopy-based screen for RhoGAP protein(s) that regulate the cortical localization of Spn1

In fission yeast, the building block of high-order Septin structures is a heterooctamer containing two molecules of each Spn1, Spn2, Spn3, and Spn4 (21). Spn1 and Spn4 are essential for the formation of the heterooctamer since high-order Septin structures are absent in cells lacking either Spn1 or Spn4 (21, 37). Therefore, to identify RhoGAP protein(s) that regulates the cortical localization of Septins, we examined the localization of Spn1, which is tagged with tdTomato (Spn1-tdTomato), in wild-type (WT) and mutant cells lacking the individual RhoGAP protein. Note that we failed to obtain the *rga1*-deletion (*rga1Δ*) strain, though nine RhoGAP proteins (i.e. Rga1-9) exist in fission yeast.

Microscopic observation showed that Spn1-tdTomato localized as scattering dots/bars on the cell cortex in interphase and appeared to be enriched at the cortical sites adjacent to the growing cell tip marked by CRIB-GFP (active Cdc42) (Figure 1A). This characteristic localization of Spn1 was found in WT and most of the RhoGAP deletion-mutant cells. Interestingly, the absence of *rga6* or *rga8* significantly impaired the characteristic localization of Spn1 on the cell cortex (Figures 1A and 1B).

**Figure 1.**
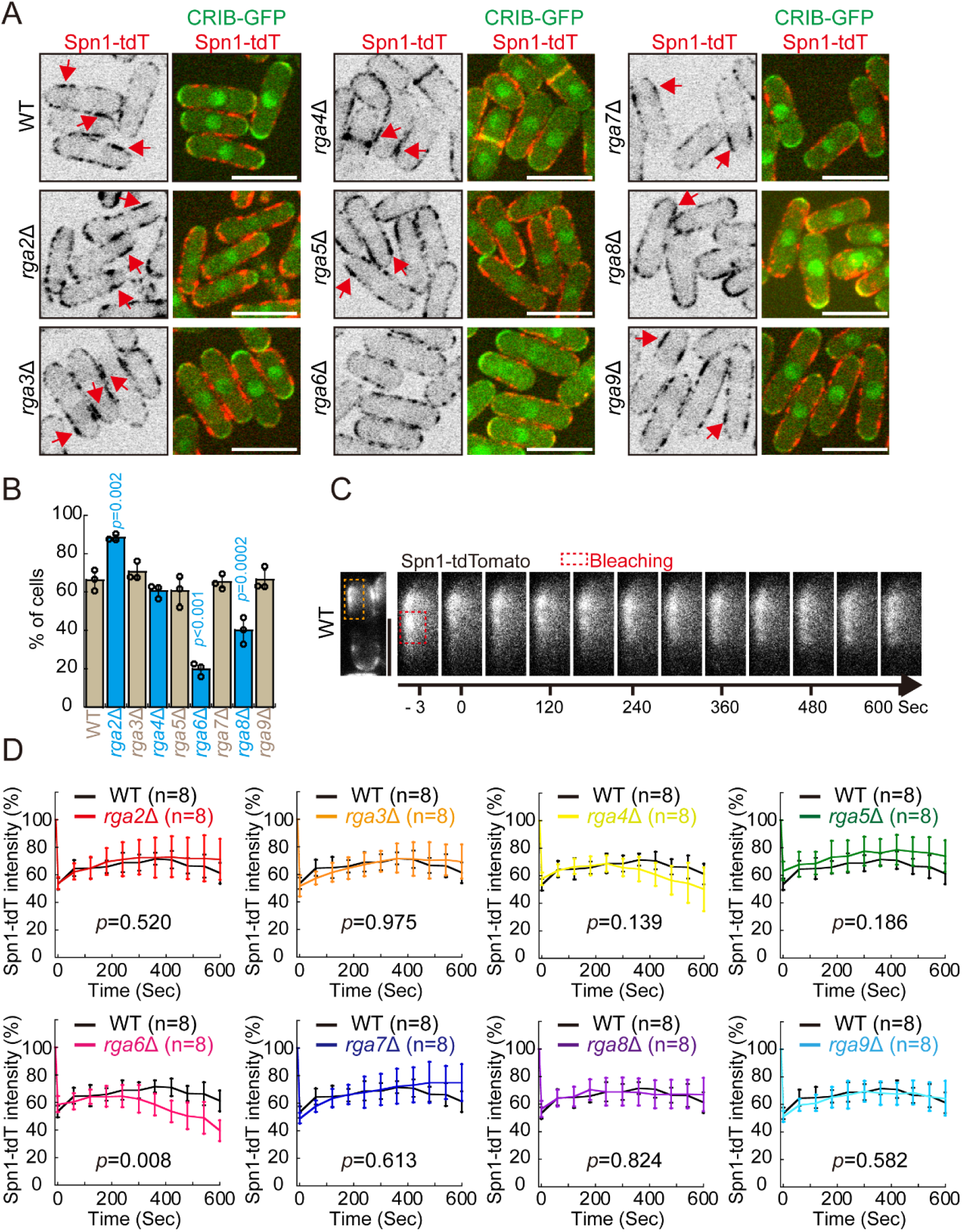
Screening for the Rho GAP proteins that are involved in regulating the cortical localization and dynamics of Spn1-tdTomato. (A) XY cross-section images of wild-type (WT) and *rga2*-deletion (*rga2*Δ), *rga3*Δ, *rga4*Δ, *rga5*Δ, *rga6Δ*, *rga7*Δ, *rga8*Δ, and *rga9*Δ cells expressing Spn1-tdTomato (Septin 1) and CRIB-GFP (indicate active Cdc42). Note that stack images consisting of 11 planes with 0.5 μm spacing were acquired but only the middle plane was shown. The red arrows indicate the accumulated Spn1 at the cortical regions near the growing cell tip, marked by CRIB-GFP. Scale bar, 10 μm. (B) Percentage of the cells displaying concentrated Spn1-tdTomato adjacent to the cell tip decorated by CRIB-GFP. Note that only cells displaying CRIB-GFP at the tip were selected for the quantification. Three independent experiments were carried out, and >60 cells were used for each quantification. One-way ANOVA analysis with Tukey HSD test was performed to calculate *p* values (vs WT). Indicated are the significant *p* values of mutant cells. (C) Fluorescence recovery after photobleaching (FRAP) experiments conducted to assess cortical Spn1 dynamics. The montage images are XY cross-section time-lapse images of the boxed region of a WT cell expressing Spn1-tdTomato. The red dashed rectangle indicates the region selected for photobleaching. Scale bar, 5 μm. (D) FRAP analysis of the recovery of Spn1-tdTomato fluorescence in WT, *rga2*Δ, *rga3*Δ, *rga4*Δ, *rga5*Δ, *rga6Δ*, *rga7*Δ, *rga8*Δ, and *rga9*Δ cells. Eight FRAP experiments were conducted for each cell type, and statistical analysis was performed by two-way ANOVA test.

We then performed fluorescence recovery after photobleaching (FRAP) experiments to assess the cortical dynamics of Spn1 in WT and the RhoGAP mutant cells. Briefly, a ~3.8 μm^2^ square region on the cell cortex near the cell tip was selected for FRAP analysis, and ~50% of the fluorescence within the selected regions was bleached. The recovery fluorescence was then monitored with a confocal scanning laser microscope. As shown in Figures 1C and 1D, the Spn1-tdTomato signals in the bleached region recovered slowly, suggesting the cortical localization of Spn1 was not very dynamic. The FRAP recovery plots of WT and the RhoGAP deletion-mutant cells except *rga6Δ* were comparable as determined by ANOVA analysis (Figure 1D). Intriguingly, in *rga6Δ* cells, the Spn1 fluorescent signals decayed over time after photobleaching, indicative of defective localization of Spn1-tdTomato to the cell cortex. Taken together, we concluded that the RhoGAP protein Rga6 plays a crucial role in regulating the cortical localization of Spn1.

### Rga6 promotes the localization of Spn1 to the cell cortex in interphase

Next, we characterized the localization of Spn1-tdTomato in interphase WT and Rga6 mutant cells. First, the intensity of Spn1-tdTomato fluorescence along the plasma membrane in WT and *rga6Δ* cells was measured by line-scan analysis (Figures 2A and 2B). The measurement results showed that Spn1-tdTomato signals along the plasma membrane were generally weaker in *rga6Δ* cells than in WT cells (Figure 2B), and the diminished signals were not due to the altered expression of Spn1-tdTomata in *rga6Δ* cells (Supplementary Figures S1A and S1B). This observation was confirmed by quantification of the average Spn1-tdTomato signal intensity along the plasma membrane (Figure 2E). We then assessed the effect of Rga6 overexpression on the cortical localization of Spn1. Microscopic analysis showed that the localization of Spn1-tdTomato to the cell cortex was remarkably enhanced in Rga6-overexpressing cells (Figures 2C, 2D and 2F), and the enhanced Spn1 signals were not due to the altered expression of Spn1-tdTomato in Rga6-GFP-overexpressing cells (supplementary Figures S1C and S1D). We further expressed Rga6-GFP from the *nmt41* promoter, a promoter capable of being repressed by thiamine, and controlled the expression levels of Rga6-GFP using 0 and 0.1 μM thiamine (Figure 2G). In the cells carrying the *nmt41* promoter, the average intensity of Rga6-GFP and Spn1-tdTomato signals along the cell cortex was measured, and correlation analysis showed that Rga6-GFP and Spn1-tdTomato signals are significantly linearly related (Figure 2G). It is possible that the enhanced cortical localization of Spn1-tdTomato in Rga6-overexpressing cells causes Spn1 to be less dynamic on the cell cortex. To test this possibility, FRAP was employed to analyze Spn1-tdTomato dynamics on the cell cortex in WT and Rga6-overexpressing cells. As shown in Figures 2H and 2I, Spn1-tdTomato signals within the bleached region in the Rga6-overexpressing cells hardly recovered, confirming that Spn1-tdTomato on the cell cortex in Rga6-overexpressing cells is almost static. These results suggest that proper localization of Septins to the cell cortex depends on Rga6.

**Figure 2.**
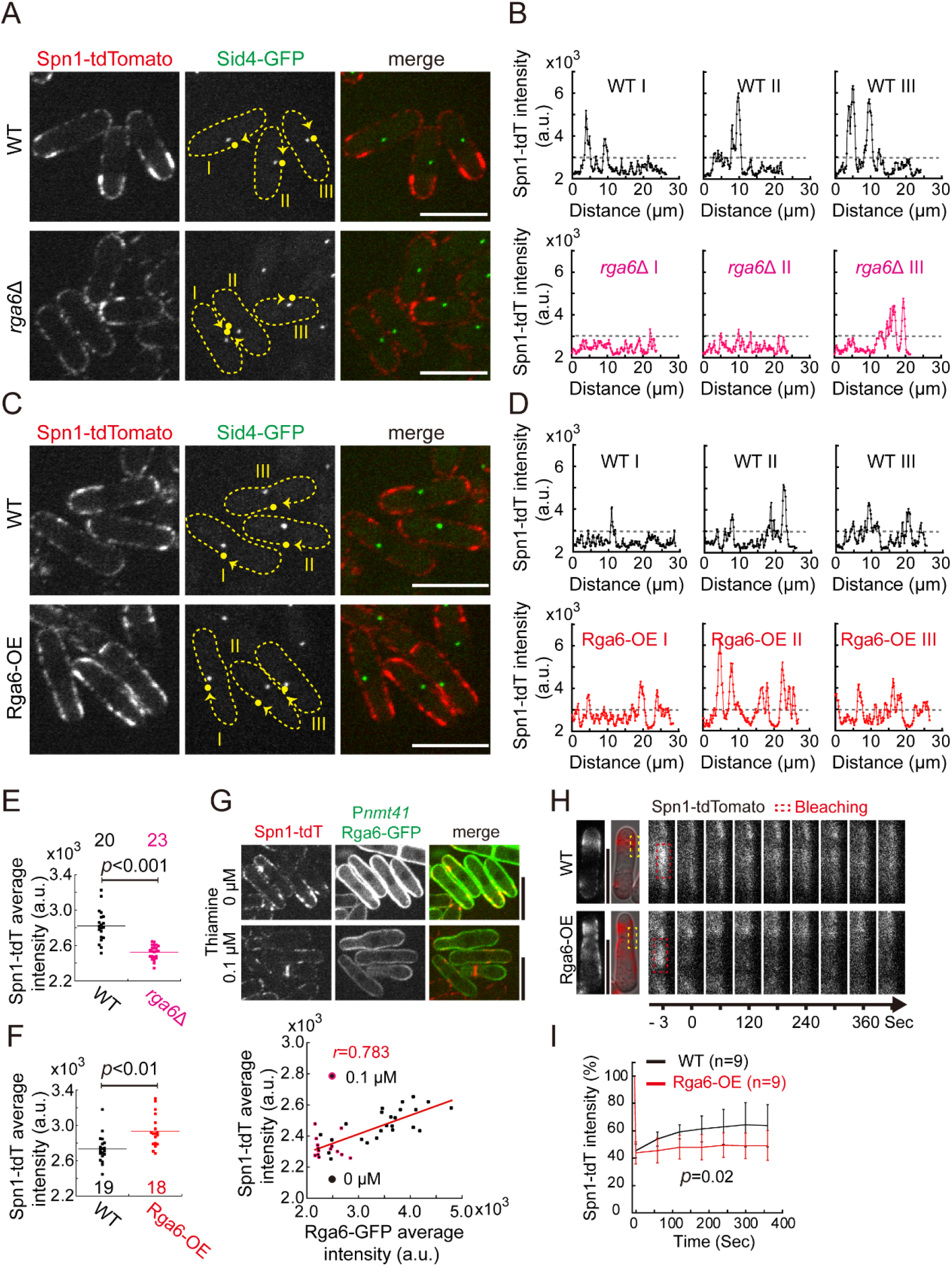
Cortical localization of Spn1-tdTomato in WT, *rga6*Δ, and Rga6-overexpressing cells. (A) XY cross-section images of WT and *rga6*Δ cells expressing Spn1-tdTomato. The cells marked with the numbered dashed lines were used for intensity measurements. Scale bar, 10 μm. (B) Line-scan analysis of Spn1-tdTomato fluorescent intensity along the cell cortex indicated in (A) by dashed lines. The filled circles are the start points. (C) XY cross-section images of WT and Rga6-overexpressing (Rga6-OE) cells expressing Spn1-tdTomato. The cells marked with the numbered dashed lines were used for intensity measurements. Scale bar, 10 μm. (D) Line-scan analysis of Spn1-tdTomato fluorescent intensity along the cell cortex indicated in (C) by dashed lines. The filled circles are the start points. (E) The average intensity of Spn1-tdTomato on the cortex of WT and *rga6Δ* cells. Student’s t-test was used to calculate the *p* value, and the number of cells analyzed is indicated. (F) The average intensity of Spn1-tdTomato on the cortex of WT and Rga6-OE cells. Student’s t-test was used to calculate the *p* value, and the number of cells analyzed is indicated. (G) Correlation analysis of the average intensity of Spn1-tdTomato and Rga6-GFP on the cell cortex. On the top panel are maximum projection images of cells expressing Spn1-tdTomato and Rga6-GFP (from the *nmt41* promoter). Scale bar, 10 μm. Cells were cultured in EMM media containing 0 or 0.1 μM thiamine. The red line on the scattering plot is a linear regression, and *r* indicates Pearson correlation coefficient; black and pink dots are data points of cells cultured in media containing 0 and 0.1 μM thiamine, respectively. (H) XY cross-section time-lapse images of the boxed region of WT and Rga6-OE cells expressing Spn1-tdTomato. The dashed red rectangles were selected for photobleaching. Scale bar, 5 μm. (I) FRAP analysis of the recovery of Spn1-tdTomato fluorescence in WT and Rga6-OE cells expressing cells. Two-way ANOVA test was used to calculate the *p* value, and the cell number analyzed is indicated.

### Rga6 physically interacts with the Septin complex

The above relationship between Spn1 and Rga6 on the cortical localization prompted us to examine colocalization of the two proteins and to test their interaction. Microscopic observation of WT cells expressing Spn1-tdTomato and Rga6-3GFP showed that Spn1-tdTomato and Rga6-3GFP colocalize on the cell cortex, particularly at the sites adjacent to the cell tip, during interphase (Figure 3A). Consistent with the published data (38), Rga6 decorated the cell sides more strongly than the cell tips during interphase. Co-immunoprecipitation assays were then performed to test the interaction between Spn1 and Rga6, Spn2 (positive control), or Bub1 (the spindle assembly checkpoint protein, serving as a negative control). As shown in Figure 3B, Spn1-tdTomato co-precipitated Spn2-3GFP and Rga6-3GFP, but not Bub1-GFP, suggesting that Spn1 interacts with Rga6 (Figure 3B). We further tested the interaction between the Septin complex and Rga6 by GST pull-down assays. Specifically, Spn1, Spn2-GST, Spn3, and Spn4 were coexpressed in *E.coli*, and the Septin complex composed of Spn1-4 (referred to as Septins) was purified with glutathione resins. We then performed pull-down experiments with GST-Septins, GST, and cell lysates containing Rga6-GFP. As shown in Figure 3C, GST-Septins, but not GST, was able to co-precipitate His-Rga6-GFP. Similarly, GST-Septins was able to co-precipitate the recombinant protein His-Rga6-GFP, but not His-GFP, and GST was not able to co-precipitate either His-Rga6-GFP or His-GFP (Figure 3D). Together, these biochemistry data suggest that Rga6 physically interacts with the Septin complex.

**Figure 3.**
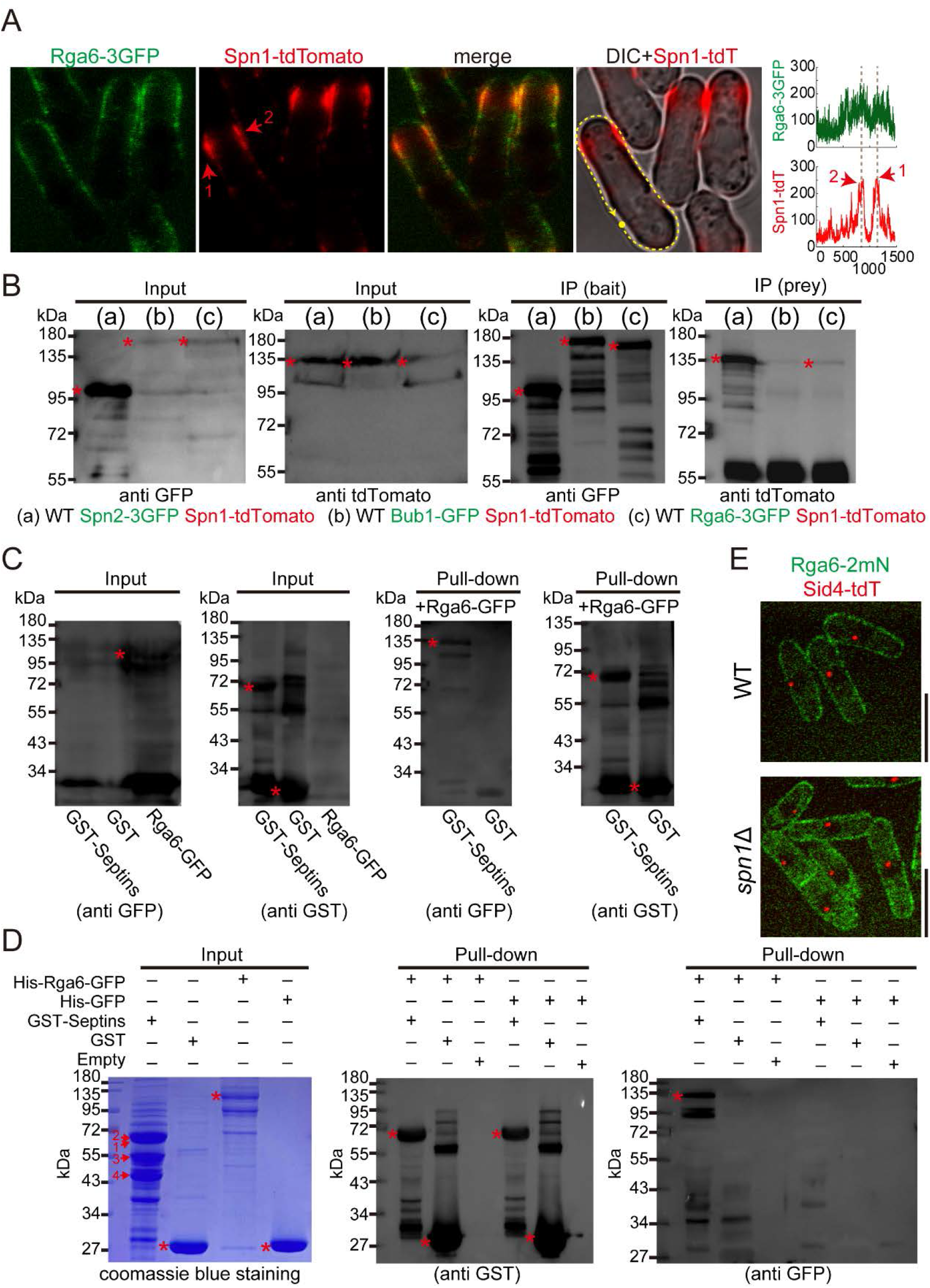
Testing colocalization and interaction between Rga6 and the septin complex. (A) Single-plane images of WT cells expressing Spn1-tdTomato and Rga6-3GFP. Note that red arrows indicate the concentrated Spn1-tdTomato near the cell tip. Linescan analysis was performed on the cell marked by the dashed line, and the average intensity of Spn1-tdTomato and Rga6-3GFP along the dashed line was plotted on the right. Scale bar, 10 μm. (B) Co-immunoprecipitation (Co-IP) experiments showing that Spn1-tdTomato interacts with Rga6-3GFP and Spn2-3GFP, but not Bub1-GFP. Note that Spn2-3GFP and Bub1-GFP were used as positive and negative controls, respectively. Asterisks indicate either GFP or tdTomato tagged proteins. Antibodies against GFP were used as baits. Western blotting was performed with antibodies against GFP and tdTomato. (C) GST pull-down assays using cell lysates containing Rga6-GFP. Asterisks indicate Rga6-GFP, Spn2-GST, or GST. Note that Spn2 in the co-purified Septin complex is fused to GST (represented by GST-Septins). Western blotting was performed with antibodies against GST and GFP. (D) GST pull-down assays with recombinant proteins. The indicated recombinant proteins were purified from *E.coli*, and GST-Septins represent the co-purified Septin complex (the numbers on the Coomassie blue staining gel indicate Spn1-4, respectively), in which Spn2 is tagged with GST. Asterisks indicate GST, Spn2-GST, His-GFP, or His-Rga6-GFP. Western blotting was performed with antibodies against GST and GFP. (E) XY cross-section images of WT and *spn1*Δ cells expressing Sid4-tdTomato and Rga6-2mNeonGreen. Scale bar, 10 μm.

We further asked whether Spn1 plays a role in localizing Rga6 to the cell cortex. Microscopic analysis showed that the localization of Rga6-2mNeonGreen on the cell cortex was comparable in WT and *spn1Δ* cells (Figure 3E), suggesting that the cortical localization of Rga6 is independent of Spn1. Hence, proper localization of Septins to the cell cortex requires Rga6, but not *vice versus*.

### Proper localization of Spn1 to the cell cortex requires the presence of cortical Rga6

Rga6p contains an N-terminal SR (Serine-rich) region (a.a., 187-253), a GAP domain (a.a., 329-547), and a C-terminal PBR domain (a.a., 700-733) (Figure 4A) (38). We attempted to identify the Rga6 domains/region(s) that interact with Spn1. To this end, co-immunoprecipitation was performed to test the interaction between Spn1 and full-length or the deletion-truncation Rga6 mutants (Figures 4A and 4B). As shown in Figure 4B, the amount of Spn1 that was co-precipitated by the truncation variants of Rga6 was much less than the amount of Spn1 that was co-precipitated by full-length Rga6. Particularly, the Rga6 variant lacking the small PBR domain, i.e. Rga6(ΔPBR), almost failed to co-precipitate Spn1 (Figure 4B). This result indicates that multiple regions in the entire protein may be required for the interaction of Rga6 with Spn1.

**Figure 4.**
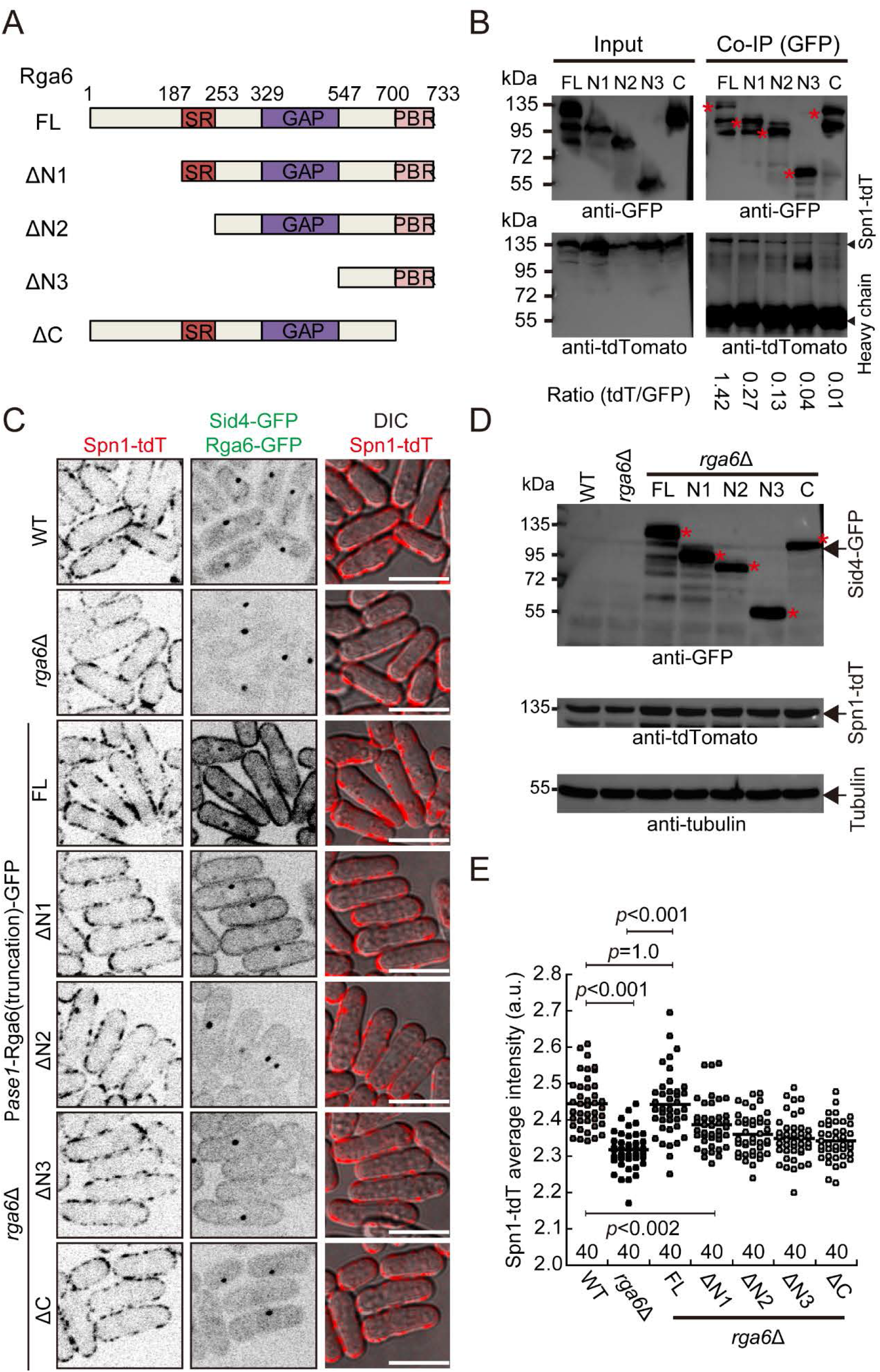
Mapping the Rga6 region(s) required for interacting with Spn1. (A) Diagram illustrating the domain structure of Rga6. Rga6 contains an N-terminal serine-rich region (SR), a middle GTPase activating protein domain (GAP), and a C-terminal polybasic region (PBR). Rga6 truncation mutants as indicated were generated for co-IP assays (B) and microscopic analysis (C). (B) Co-IP assays testing the interaction between Spn1 and the Rga6 truncation mutants. The strains indicated in (C) were used for the co-IP assays, and GFP-fusion proteins served as baits. The band intensity ratio of tdTomato over GFP was indicated at the bottom of each lane. (C) XY cross-section images of *rga6*Δ cells expressing Spn1-tdTomato and the indicated Rga6-GFP variants from the *ase1* promoter. Scale bar, 10 μm. (D) Testing expression of Spn1-tdTomato and the GFP-fused proteins in the strains indicated in (C). Western blotting analysis was performed using antibodies against GFP, tdTomato, and tubulin. Asterisks mark GFP-fused proteins. (E) Dot plot of the average intensity of cortical Spn1-tdTomato in *rga6Δ* cells expressing the indicated Rga6-GFP variants in (C). One-way ANOVA test was used to calculate the *p* values (vs FL), and the cell number analyzed is indicated.

The expression of Rga6 from its own promoter appeared to be very weak (Figure 3E). Therefore, full-length Rga6 and its truncation variants tagged with GFP were expressed from the *ase1* promoter for microscopic observation (Figure 4C). Expression of the GFP-tagged proteins was confirmed by western blotting analysis (Figure 4D), and the expression levels of Spn1-tdTomato did not appear to be altered significantly (Figure 4D). Interestingly, Rga6(ΔN2)-GFP and Rga6(ΔN3)-GFP signals on the cell cortex were very weak and Rga6(ΔPBR)-GFP did not localize to the cell cortex. By contrast, the cortical signals of Rga6(FL)-GFP and Rga6(ΔN1)-GFP were apparent. Quantification of the cortical localization of Spn1-tdTomato showed that the average fluorescent intensity of Spn1-tdTomato along the cell cortex was fully rescued by ectopic expression of full-length Rga6-GFP and, to a lesser degree, by Rga6(ΔN1)-GFP in *rga6*Δ cells (Figure 4E). Rga6(ΔN2)-GFP, Rga6(ΔN3)-GFP, and Rga6(ΔPBR)-GFP did not appear to significantly rescue the localization of Spn1-tdTomato in *rga6*Δ cells (Figure 4E). Collectively, these results suggest that both the presence of Rga6 on the cell cortex and the multiple domains/regions at the N-terminus of Rga6 are required for directing the proper interaction between Rga6 and Spn1 and the localization of Spn1 to the cell cortex.

### The presence of both Spn2 and Rga6 enhances their localization to reconstituted liposome membranes

We then sought to test the membrane-binding affinity of the recombinant proteins Rga6 and Septins by *in vitro* liposome reconstitution assays. Using Phosphatidylcholines (PC) and Phosphatidylserines (PS), we followed the previously developed emulsion-based method to generate single layer liposomes enclosing recombinant proteins (36), and the protein-enclosing liposomes were then imaged with a spinning-disk microscope (Figure 5A). Among the Septin complex, Spn2 alone was able to be expressed and purified from *E.coli*. Therefore, we tagged Spn2 with BFP and purified recombinant proteins Spn2-BFP and BFP, respectively, from *E.coli*. Rga6-GFP was expressed in insect cells. All recombinant proteins except Rga6-GFP were further purified by size-exclusion chromatography (Figure 5B). First, we examined the localization of each recombinant protein alone. As shown in Figure 5C, in most of the Rga6-GFP-enclosing liposomes (~74%), Rga6-GFP diffused within the capsule and also displayed faint cortical localization. Similarly, BFP diffused within ~76% of BFP-enclosing liposomes. Spn2-BFP displayed a single focus and regional localization on the cortex of ~64% and ~36% of Spn2-BFP-enclosing liposomes, respectively. Remarkably, the presence of both Rga6-GFP and Spn2-BFP enhanced the localization of both proteins to the cortex of liposomes since Rga6-GFP and Spn2-BFP showed colocalization on the cortex of ~80% of liposomes (Figure 5D). By contrast, in ~83% of liposomes, Rga6-GFP localized to the cortex but no BFP was found on the cortex of the liposomes (Figure 5D). Together, these results support the conclusion that Rga6 interacts with Septins and promotes the localization of Septins to the cortical membrane.

**Figure 5.**
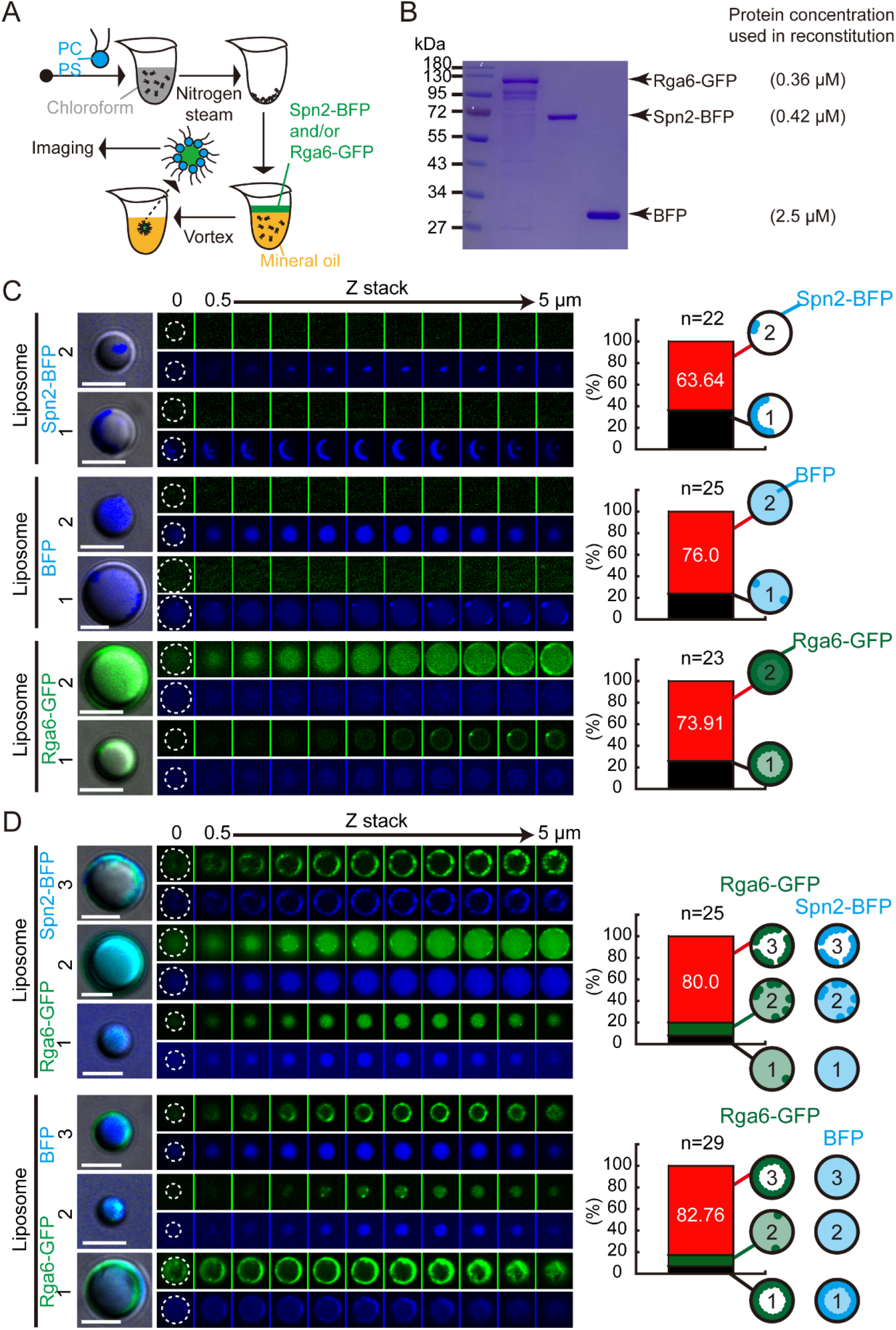
Liposome reconstitution to test the membrane-binding ability of Spn2 and Rga6. (A) Diagram illustrating the procedure used for the liposome reconstitution. In the lipid mixture, 90% Phosphatidylcholine (PC) and 10% Phosphatidylserine (PS) were used, and single-layer liposomes enclosing Spn2-BFP and/or Rga6-GFP were generated with mineral oil and were imaged with a spinning-disk microscope. (B) Recombinant proteins used in the liposome reconstitution assay were analyzed by Coomassie blue staining. Note that the protein concentration in the reconstitution experiments is shown on the right. (C) Representative Z-stack images of liposomes enclosing Spn2-BFP, BFP, or Rga6-GFP, respectively. Dashed lines mark liposome outlines. Two types of protein localization patterns were shown. Spn2-BFP: 1) regional localization on the liposome (36.36%) and 2) one focus on the liposome (63.64%). BFP: 1) diffuse within the liposome but with a few foci on the liposome (24%) and 2) diffuse within liposome (76%). Rga6-GFP: 1) mainly localize to the liposome but with a few foci on the liposome (26.09%) and 2) localize to the liposome and also diffuse within the liposome (73.91%). Quantification and diagrams illustrating the localization patterns of the recombinant proteins are shown on the right. Scale bar, 5 μm. (D) Representative Z-stack images of liposomes enclosing the mixture of Rga6-GFP and Spn2-BFP or Rga6-GFP and Spn2-BFP. Dashed lines mark liposome outlines. Three types of protein localization patterns were found. Rga6-GFP plus Spn2-BFP: 1) Rga6-GFP and Spn2-BFP diffuse within the liposome (8%), 2) Rga6-GFP and Spn2-BFP colocalize on the liposome and also diffuse within the liposome (12%), and 3) Rga6-GFP and Spn2-BFP colocalize on the liposome (80%). Rga6-GFP plus BFP: 1) Rga6-GFP and BFP colocalize on the liposome (6.90%), 2) Rga6-GFP and BFP diffuse within the liposome but a few Rga6-GFP foci are present on the liposome (10.34%), and 3) Rga6-GFP localizes to the liposome whereas BFP diffuses within the liposome (82.76%). Quantification and diagrams illustrating the localization patterns of the recombinant proteins are shown on the right. Scale bar, 5 μm.

### Septins and Rga6 synergize to confine active Cdc42 to the growing cell tip

What is the function of the colocalized Rga6 and Spn1 on the cell cortex? It has been demonstrated that Rga6 is a GAP for Cdc42 (29). Therefore, it is conceivable that Spn1 and Rga6 may work in concert to regulate cell polarity. To test this hypothesis, we examined the localization of Spn1 and active Cdc42 (marked by CRIB-GFP) simultaneously in WT, *rga6Δ*, and Rga6-overexpressing (Rga6-OE) cells by spinning-disk live-cell microscopy. As shown in Figure 6A, in WT cells, Spn1 was excluded from the growing cell tip where active Cdc42 localized but was enriched at the adjacent regions to the growing cell tip. This was confirmed by line-scan analysis of the fluorescent intensity of Spn1-tdTomato and CRIB-GFP along the cell cortex (Figure 6B and supplementary Figure S2). By contrast, in *rga6*Δ cells, no obvious regional enrichment of Spn1-tdTomato was detected on the cell cortex while active Cdc42 appeared to spread over a wider area at the growing cell tip (Figures 6A and 6B). In sharp contrast to *rga6*Δ cells, Rga6-OE cells displayed very dense Spn1-tdTomato on the cell cortex except at the growing cell tip and active Cdc42 appeared to be restricted to a much smaller area (Figures 6A and 6B). We went on to measure the length of CRIB-GFP at the growing cell tips in WT, *rga6Δ*, Rga6-OE, and *spn1Δ* Rga6-OE cells (Figure 6C). Consistent with the live-cell imaging data (Figure 6A), the CRIB-GFP length was slightly but significantly longer in *rga6Δ* cells than in WT cells, but significantly shorter in Rga6-OE cells than in WT cells (Figure 6D). Considering the canonical role of Septins in diffusion restriction (10), we interpreted these data as evidence supporting a model that the Rga6-dependent localization of Septins to the cell cortex functions to confine active Cdc42 to the growing cell tip. Indeed, the absence of Spn1 was able to lengthen CRIB-GFP at the cell tips in Rga6-OE cells (Figures 6C and 6D). Taken together, it is highly likely that Rga6 promotes the accumulation of Septins on the cortical sites near the growing cell tip to confine active Cdc42 for regulating cell growth.

**Figure 6.**
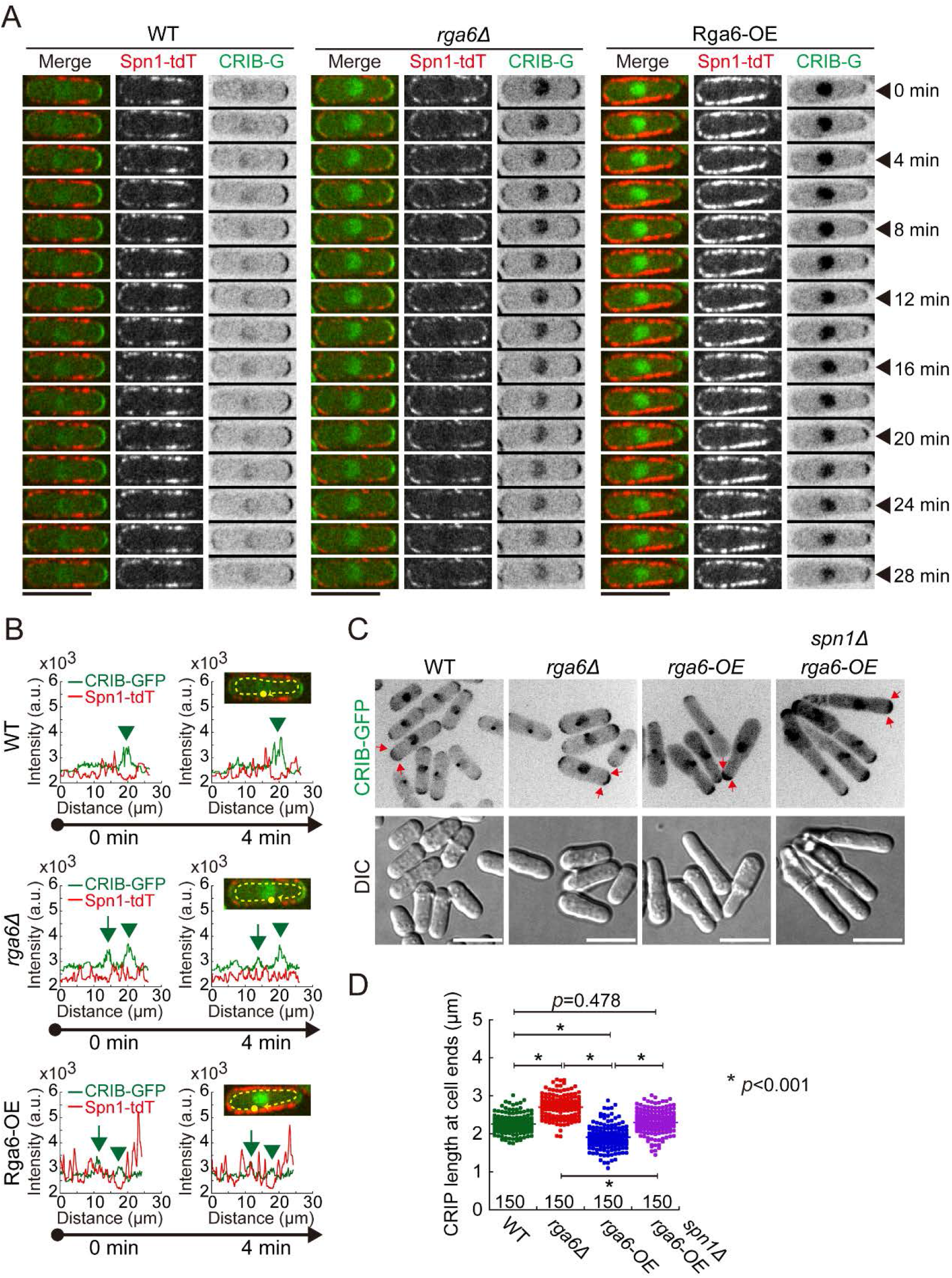
Localization of Spn1 to the cell cortex is required for directing proper localization of active Cdc42 at the cell tip. (A) XY cross-section time-lapse images of WT, *rga6Δ*, and Rga6-OE cells expressing Spn1-tdTomato and CRIB-GFP (marking active Cdc42). Scale bar, 10 μm. (B) Line-scan fluorescent intensity analysis of CRIB-GFP and Spn1-tdTomato along the dashed lines for the cells indicated in (A) at 0 min and 4 min, respectively. (C) Maximum projection images of WT, *rga6Δ*, Rga6-OE (from the *nmt41* promoter in WT and in *spn1Δ* cells, respectively) cells expressing CRIB-GFP. Red arrows mark the two edges of the crescent CRIB-GFP shapes at cell tips. Scale bar, 10 μm. (D) CRIB-GFP length at cell tips in the indicated cells. *p* values were calculated by One-way ANOVA analysis with Tukey HSD test, and the total number of cells from three independent experiments are indicated.

## DISCUSSION

Cortical localization of Septins allows assembly of Septin high-order structures required for directing a wide range of crucial cellular activities, including membrane compartmentalization, membrane remodeling, and cytokinesis (1, 3, 4). How the cortical localization of the Septin complex is regulated in space and time is, therefore, a fundamental question to be addressed. In this present study, we identified the RhoGAP Rga6 as an interacting protein of the Septin complex (Figure 3) and demonstrated that Rga6 is required for the proper localization of Septins to the cell cortex (Figure 2). Our data support a model that Rga6 promotes the localization of Septins to the cell cortex, particularly to the region near the growing cell tip, by which the GAP activity of Rga6 and the diffusion-barrier function of Septins may be integrated to regulate polarized localization of active Cdc42 (Figure 7).

**Figure 7.**
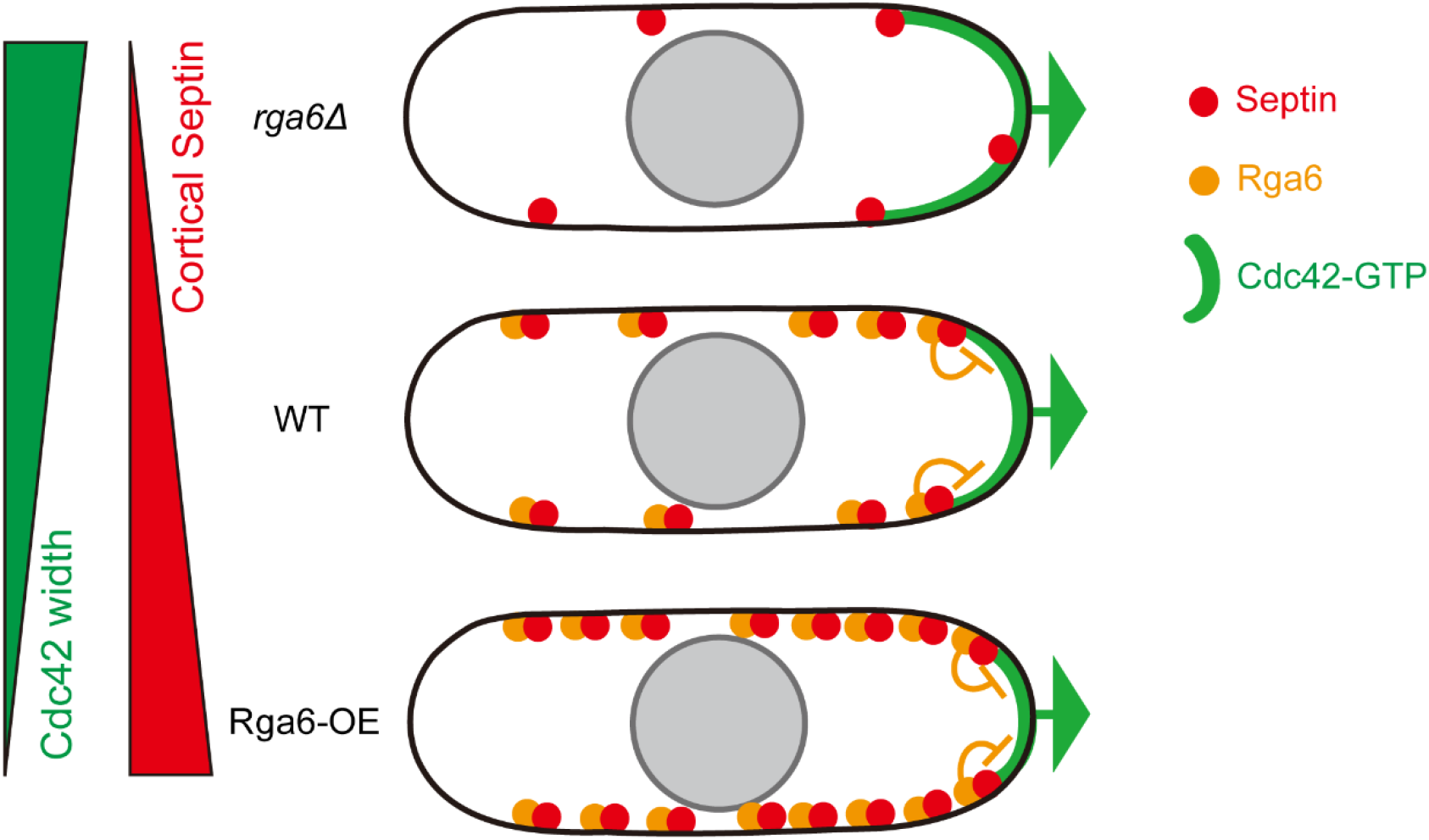
The synergistic role of Rga6 and Septin in regulating polarized cell growth. In wild type cells, Rga6 promotes the localization of Septin to the cortical region adjacent to the growing cell tip, by which polarized growth is confined and maintained. The absence of Rga6 impairs the enrichment of Septin near the growth, thus lengthening cell width. By contrast, overexpression of Rga6 abnormally enhances the cortical enrichment of Septin, leading to shortened cell width.

It is proposed that the polybasic stretch at the N-terminus enables Septins to bind negatively charged lipids on the membranes (3, 17, 18). Particularly, *in vitro* biochemical work establishes that Septins have an affinity for phosphoinositides (6, 17, 18, 20), and membrane association appears to promote assembly of Septin high-order structures (6). Similarly, we found that Spn2 alone is sufficient to bind the liposome membranes reconstituted with PC and PS though the cortical localization of Spn2 appears to be weak (Figures 5C). This finding is consistent with the general view that the cortical localization of Septins is due to their intrinsic membrane binding ability. In this regard, Septins should decorate the entire cell cortex homogenously. However, within the fission yeast cells, Septin enrichment is often found near the growing cell tip (Figures 2A and 6A). One possible explanation is that the cortical Septin localization is also regulated by assisting proteins. In this present study, we found that the presence of Rga6, a RhoGAP protein for Cdc42, greatly enhances the ability of Septins in binding liposome membranes (Figures 5C and 5D). The enhancement effect is also true within the cell because the absence of Rga6 and Rga6 overexpression compromises and enhances the cortical localization of Spn1, respectively (Figure 2). Moreover, Rga6 interacts with the Septin complex and colocolizes with the Septin complex on the cell cortex (Figure 3). Therefore, our work establishes that in addition to the intrinsic membrane binding ability, Septins depend on the RhoGAP Rga6 for localizing to the cell cortex.

Rga6 is a multi-domain protein containing a polybasic region at the extreme C-terminus, and the polybasic region is essential for localizing Rga6 to the cell cortex (Revilla-Guarinos et al., 2016) (also see Figure 4C). Although Rga6 is required for the proper localization of Septins to the cell cortex, the absence of Spn1 does not appear to affect the localization of Rga6 (Figure 3E). Therefore, we favor the model that Rga6 localizes to the cell cortex through its C-terminal polybasic region and subsequently recruits Septins to the cell cortex. It is likely that multiple regions/domains in Rga6 are responsible for interacting with Septins because deletion of the N-terminal regions in Rga6 compromises the interaction between the Rga6 mutants and Spn1 and the localization of Septins to the cell cortex (Figures 4B and 4C). It has been established that membrane interaction can promote Septin assembly (6) and Septins generally form low-order structures within the cytoplasm (3, 5). Therefore, it is conceivable that the involvement of Rga6 in localizing Septins to the cell cortex can locally concentrate low-order Septin complexes from the cytoplasm and subsequently promote formation of high-order Septin structures on the cortex.

Rga6 is a highly dynamic molecule on the cell cortex (38), and yet Septin filaments on the cell cortex are less dynamic (Figures 1D and 2H). How could two molecules with distinctly different dynamics on the cell cortex interact with one another? One possible explanation is that the interaction between the two molecules is transient. Consistently, the interaction between Spn1 and Rga6 is much weaker than the interaction between Spn1 and Spn2 (Figure 3B). The transient interaction may allow local enrichment of Septins, by a collective effort of many Rga6 molecules, and the enrichment then promotes assembly of Septin filaments. This awaits further biochemical investigation.

Septins are generally enriched on the cortical region adjacent to the growing cell tip (Figures 1A, 2A, and 6A). Our findings also revealed that Rga6 expression levels affect both the cortical localization of Septins and the tip localization of active Cdc42 (Figures 6A and 6B). Specifically, the absence of Rga6 compromises the cortical localization of Spn1 and enables active Cdc42 to spread over a wider area at the cell tip, while Rga6 overexpression enhances the cortical localization of Spn1 and restricts active Cdc42 to a smaller area at the cell tip (Figure 6). Importantly, the restriction effect of Rga6-overexpression on Cdc42 at the cell tip depends on Spn1 (Figures 6C and 6C). Considering the canonical function of Septin filements and Rga6 as a diffusion barrier and a Cdc42 GAP, respectively, we favor a model that the confinement of active Cdc42 to the growing cell tip may be regulated by a synergistic effect of Septins and Rga6, i.e. Septin-dependent compartmentalization and Rga6-dependent Cdc42 GAP activity.

The tight relationship between Septins and Cdc42 has been established previously in *Saccharomyces cerevisiae* (23–25). Instead of Cdc42 GAP proteins, the Cdc42 effectors Gic1 and Gic2 play an important role in recruiting Septins to the budding site (25). Interestingly, at the budding site, Septins also interact with Cdc42 GAP proteins and recruit the GAP proteins to inactivate Cdc42 (26). Therefore, the interaction between Cdc42 GAP proteins and Septins may have been conserved through evolution but evolves to function differently in an organism-dependent manner. In human cells, similar to Rga6 in this case, many of the RhoGAPs, particularly the Cdc42 GAP proteins, contain membrane-binding domains (27). Given the conservative nature of Septins, it would be interesting to explore whether the RhoGAPs containing membrane-binding domains also similarly regulate the cortical localization and functions of Septins in higher eukaryotic cells.

## ABBREVIATIONS

FRAP: Fluorescence recovery after photobleaching
RhoGAP: GTPase-activator protein for Rho-like GTPases

## ACKNOWLEDGEMENTS

We thank Dr. Erfei Bi (UPENN) for suggestions and members in the Fu laboratory for insightful discussion. This work is supported by grants from National Key Research and Development Program of China (2017YFA0503600), National Natural Science Foundation of China (32070707, 31871350, and 31621002), the Strategic Priority Research Program of the Chinese Academy of Sciences (XDB19040101).

## AUTHOR CONTRIBUTIONS

C.F. conceived the project. S.Z., B.Z., Z.L., and W.W. performed experiments. S.Z. analyzed data. S.Z., W.W., and C.F. wrote the paper. C.F. supervised the work. All authors made comments.

## Supplementary Information for

**Supplementary Figure S1.**
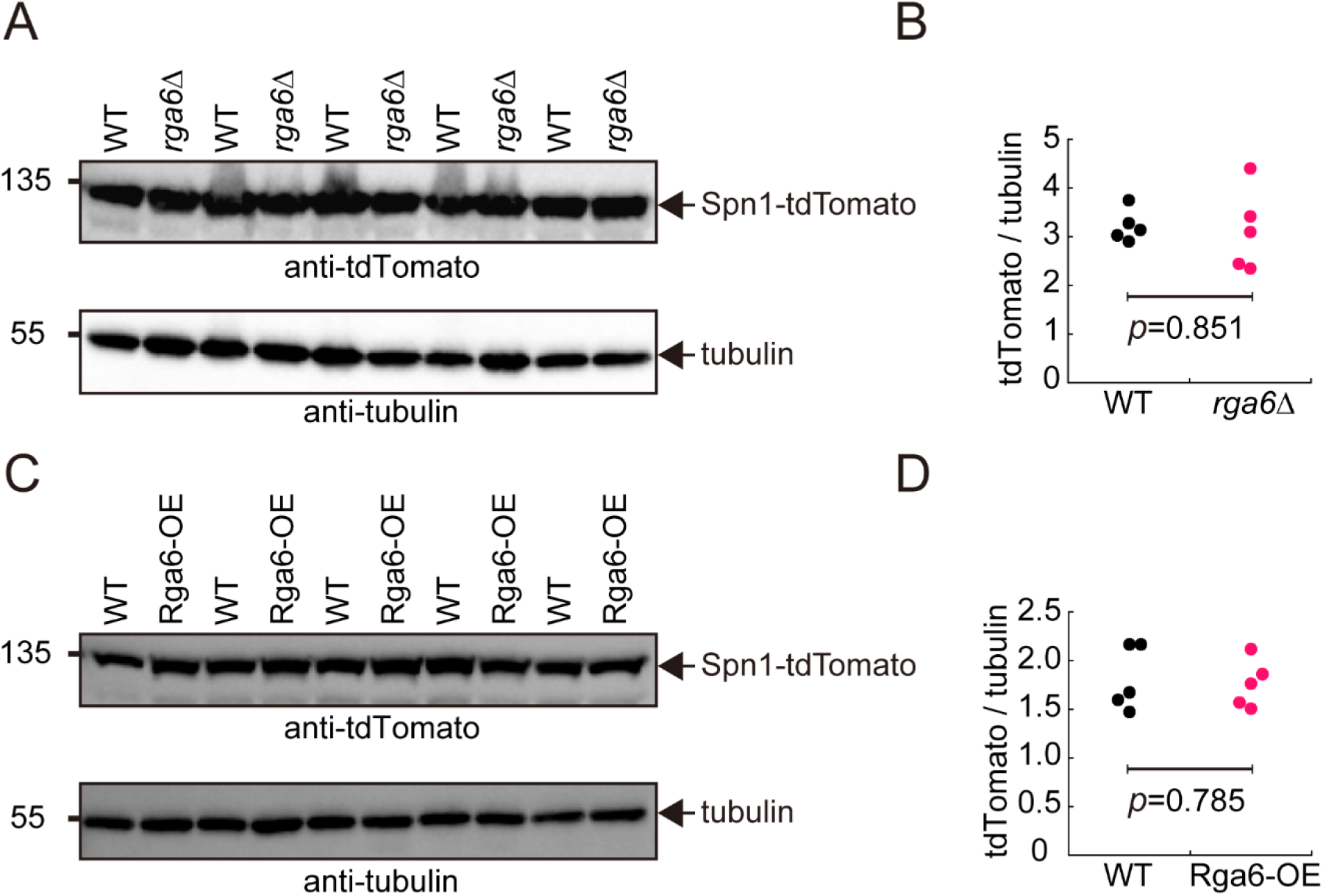
Testing the expression of Spn1-tdTomato (Related to main Fig. 2). (A and B) Western blotting analysis of Spn1-tdTomato in WT and *rga6*Δ cells. Five cultures for WT and *rga6*Δ cells were collected for analysis Antibodies against tdTomato and tubulin were used. Intensity ratio of tdTomato over tubulin is shown in (B). Statistical analysis was performed by student’s t-test. (C and D) Western blotting analysis of Spn1-tdTomato in WT and Rga6-overexpressing cells. Five cultures for WT and Rga6-overexpressing cells were collected for analysis. Antibodies against tdTomato and tubulin were used. Intensity ratio of tdTomato over tubulin is shown in (D). Statistical analysis was performed by student’s t-test.

**Supplementary Figure S2.**
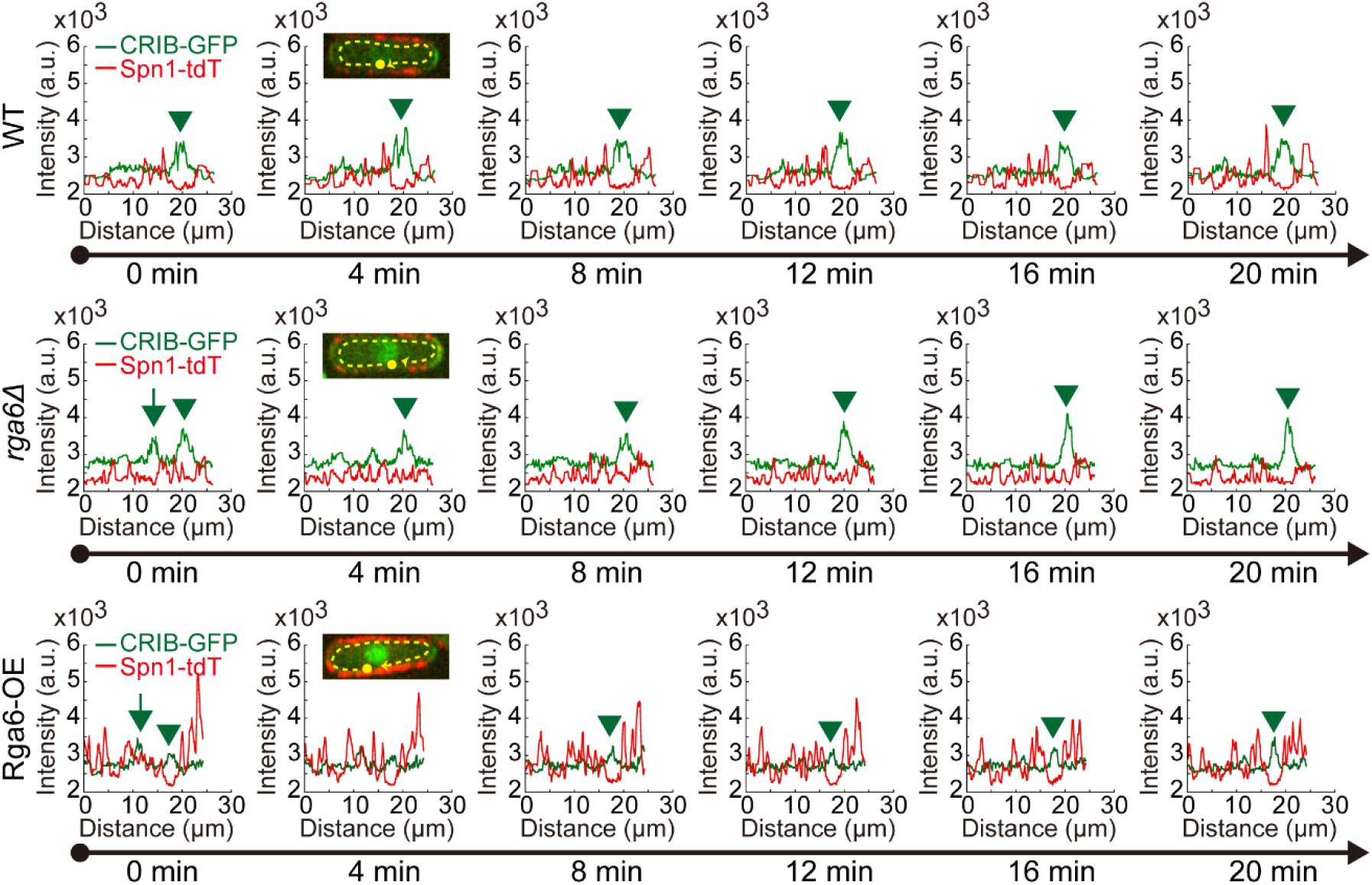
(Related to main Fig. 6). Line-scan fluorescent intensity analysis of CRIB-GFP and Spn1-tdTomato along the dashed lines for the cells indicated in (main Figure 6A) at the indicated timepoints.

## Supplementary Tables

**Table S1.**
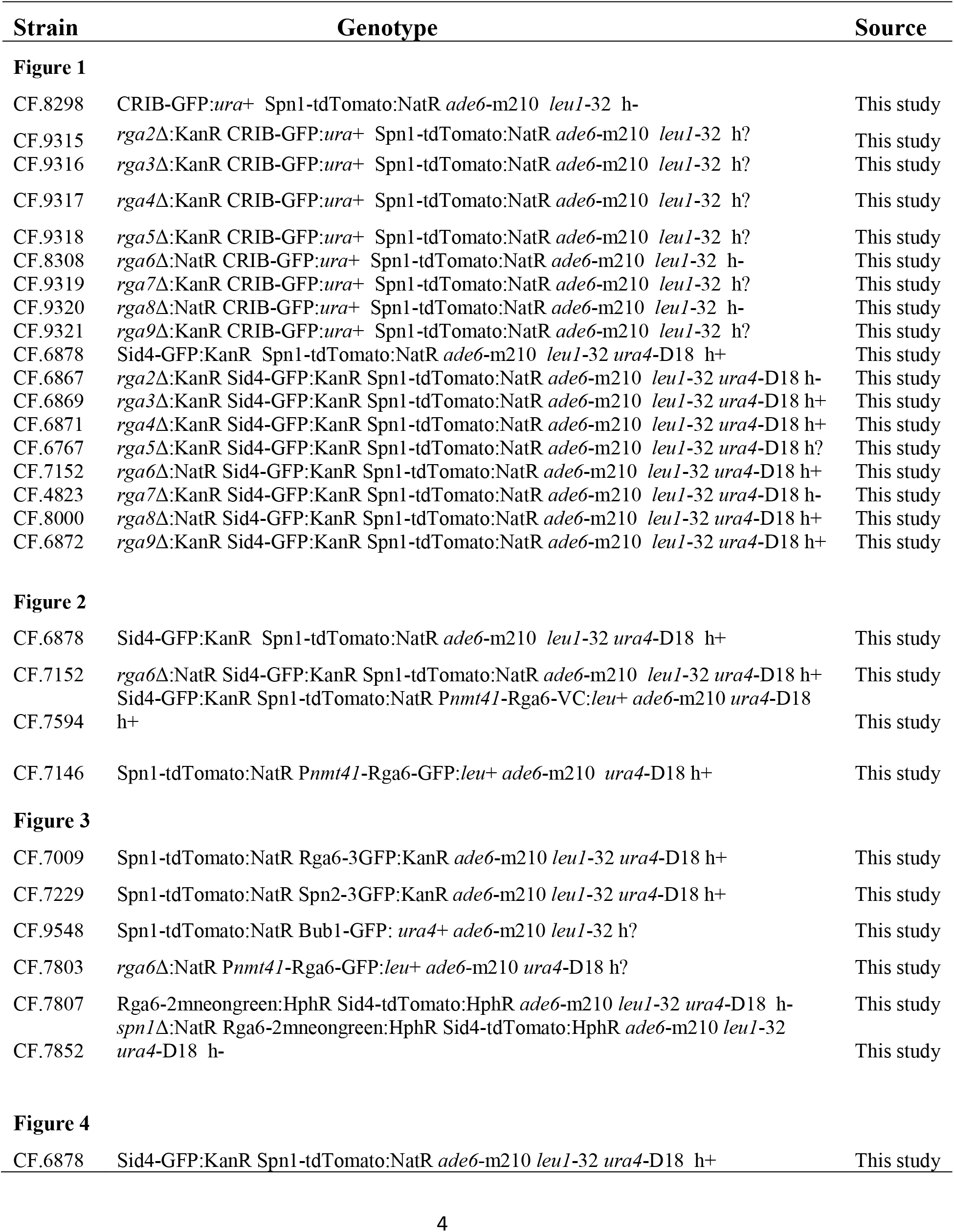
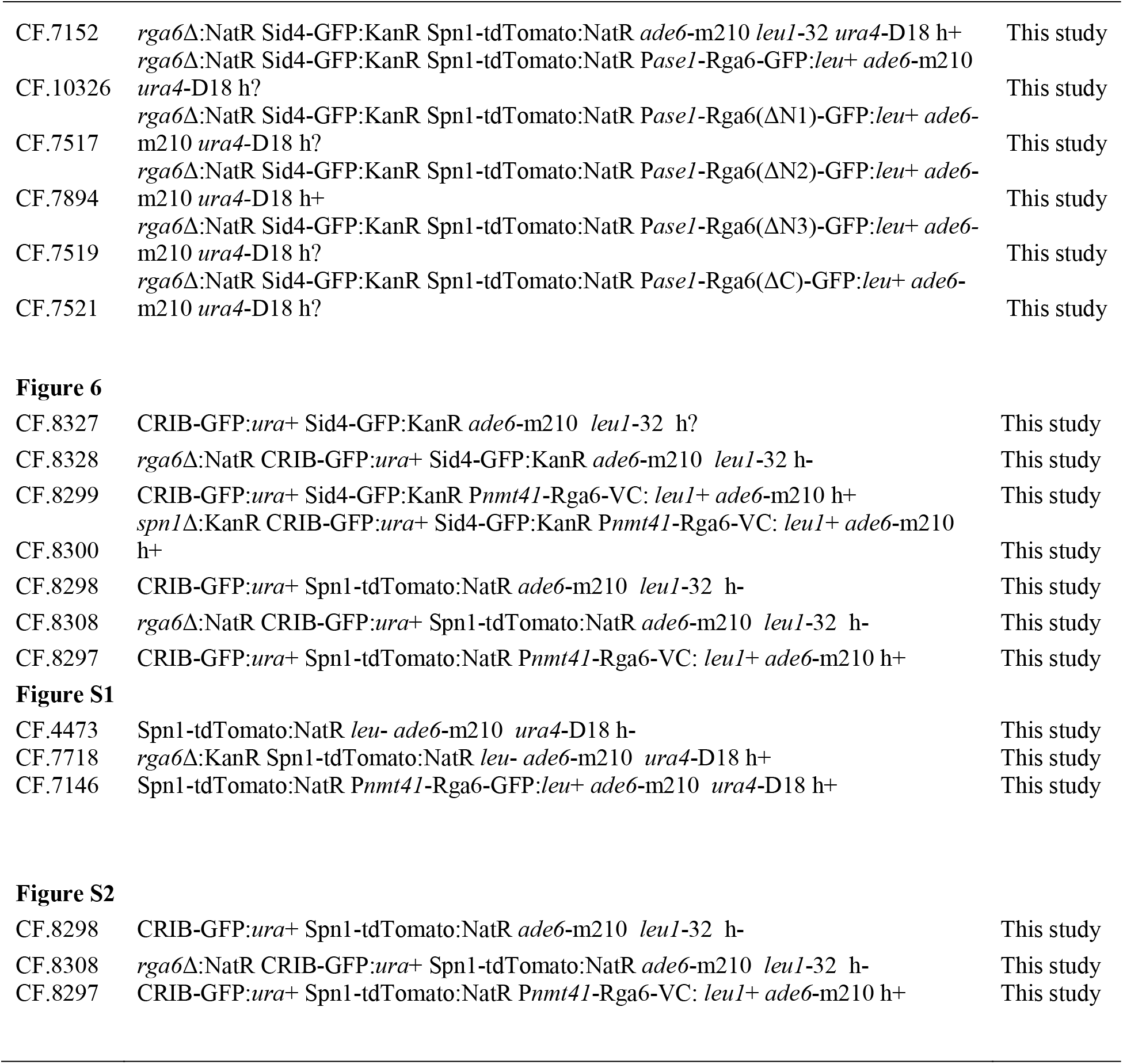
Yeast strains

**Table S2.**
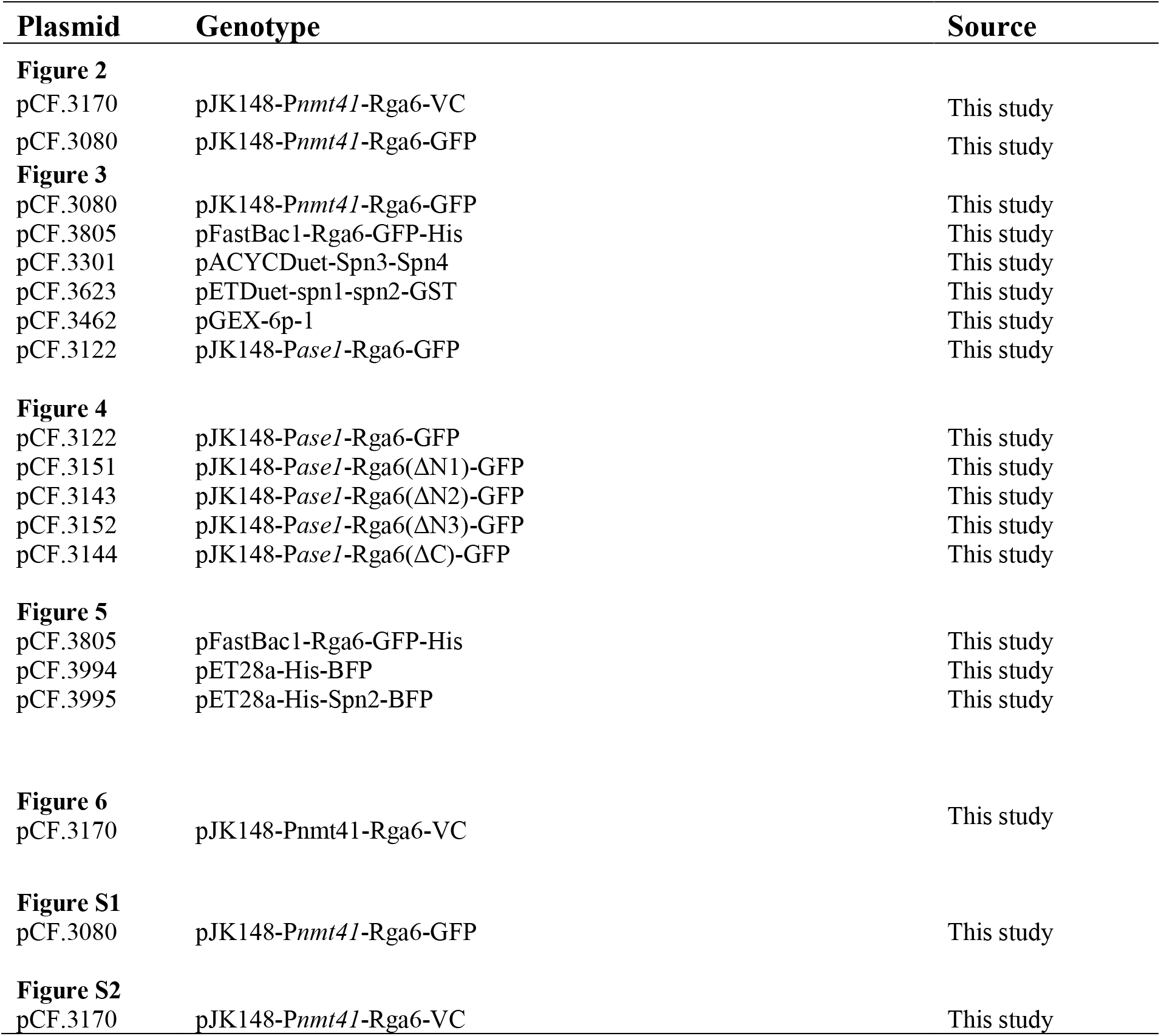
Plasmids

| **Plasmid** | **Genotype** | **Source** |
| --- | --- | --- |
| **Figure 2** |  |  |
| pCF.3170 | pJK148-*Pnmt41*-Rga6-VC | This study |
| pCF.3080 | pJK148-*Pnmt41*-Rga6-GFP | This study |
| **Figure 3** |  |  |
| pCF.3080 | pJK148-*Pnmt41*-Rga6-GFP | This study |
| pCF.3805 | pFastBac1-Rga6-GFP-His | This study |
| pCF.3301 | pACYCDuet-Spn3-Spn4 | This study |
| pCF.3623 | pETDuet-spn1-spn2-GST | This study |
| pCF.3462 | pGEX-6p-1 | This study |
| pCF.3122 | pJK148-*Pase1*-Rga6-GFP | This study |
| **Figure 4** |  |  |
| pCF.3122 | pJK148-*Pase1*-Rga6-GFP | This study |
| pCF.3151 | pJK148-*Pase1*-Rga6(ΔN1)-GFP | This study |
| pCF.3143 | pJK148-*Pase1*-Rga6(ΔN2)-GFP | This study |
| pCF.3152 | pJK148-*Pase1*-Rga6(ΔN3)-GFP | This study |
| pCF.3144 | pJK148-*Pase1*-Rga6(ΔC)-GFP | This study |
| **Figure 5** |  |  |
| pCF.3805 | pFastBac1-Rga6-GFP-His | This study |
| pCF.3994 | pET28a-His-BFP | This study |
| pCF.3995 | pET28a-His-Spn2-BFP | This study |
| **Figure 6** |  |  |
| pCF.3170 | pJK148-*Pnmt41*-Rga6-VC | This study |
| **Figure S1** |  |  |
| pCF.3080 | pJK148-*Pnmt41*-Rga6-GFP | This study |
| **Figure S2** |  |  |
| pCF.3170 | pJK148-*Pnmt41*-Rga6-VC | This study |

